# Structurally Informed Fitness Landscapes for Surveillance of Emerging PRRSV Variants

**DOI:** 10.1101/2025.10.06.680729

**Authors:** Vaishnavey SR, Supantha Dey, Sakib Ferdous, Riza Danurdoro, Michael A Zeller, Ratul Chowdhury

**Author notes:** Present address: 6756 Gables Way, Johnston, Iowa, USA 50131.

## Abstract

Antibodies play a central role in neutralizing pathogens through direct interference with viral entry and recruitment of effector immune cells. However, viruses employ sophisticated escape mechanisms to evade these defenses, primarily through mutations in surface glycoproteins that reduce antibody binding affinity or alter critical functional domains. Antibody escape remains a formidable challenge in drug design efforts for Porcine Reproductive and Respiratory Syndrome Virus (PRRSV), a pathogen notorious for its rapid evolution and structural plasticity. Here we present EscaPRRS (Escape scoring for PRRS virus), a Bayesian variational autoencoder trained on ESM-2 protein language model embeddings to predict the fitness landscape and escape propensities of 52,622 mutants (recorded over 10 years) of the immunodominant GP5 glycoprotein encoded by PRRSV *ORF5* gene. Unlike conventional models that estimate escape propensity only from sequence information, EscaPRRS circumvents the need for extensive alignments, integrating contributions from surface accessibility and biochemical dissimilarity at the binding interface. Our escape propensity scores demonstrate reliable structural and biological fidelity, with EscaPRRS scores correlating with binding affinities on seven different porcine receptor proteins (Pearson r = 0.74). Notably, EscaPRRS captures seasonal trends in immune evasion, highlighting its applicability in forecasting and surveillance of emerging/re-emerging PRRSV variants.

**IMPORTANCE:** Porcine Reproductive and Respiratory Syndrome Virus (PRRSV) remains the most economically detrimental illness for swine products, causing more than $1.2 billion in annual production loss in the United States, with indirect implications to food security and human health. PRRSV infection is known to severely affect porcine alveolar macrophages (PAMs), causing respiratory difficulties, following blockage of inflammatory signals that aids easy viral reproduction in the host cell. Identifying critical amino acid mutations at the highly variant GP5 glycoprotein attached to the viral cell membrane provides information on structural features linked to antigenic diversity and antibody neutralization, that potentially lead to clinical outbreaks in sow farms. Our effort employs machine learning approaches to learn patterns from a large number of sequences to map these critical domains to three-dimensional structures to score them for antibody escape tendencies. This greatly enhances our understanding of receptor and antibody binding mechanisms in PRRSV GP5.

## INTRODUCTION

Porcine Reproductive and Respiratory Syndrome Virus (PRRSV) is a rapidly evolving pathogen with complex structural and immune evasion mechanisms, characterized by high frequency mutations and recombination between different lineages^1^, reported to have the fastest evolutionary rate of all known RNA viruses.

This evolutionary agility challenges and eludes conventional vaccine design paradigms, particularly directed evolution and rational design, which, despite their historical contributions, struggle to keep pace with viral plasticity. The inherent bias in directed evolution towards selecting variants that confer short-term immune escape suggests a semi-rational design approach that accounts for the dynamic and disordered regions of viral proteins^2^.

Contemporary PRRSV is divided into PRRSV Europensis (PRRSV-1) and PRRSV Americanese (PRRSV-2), which are colloquially referred to as the European PRRSV and American PRRSV respectively (ICTV). These two genotypes are recognized to have diverged by at least 1980, though their divergence may have occurred earlier^3^. While PRRSV-1 is a noted disease within North America, it is in much lower circulation making up no more than 1% of PRRSV detections per year since 2015 based on aggregate data provided by the major swine diagnostic laboratories who contribute to the Swine Disease Reporting System^4^. PRRSV-2 in contrast, has composed more than 99% of PRRSV detections in the United States over the same time period and has contributed to significant disease and mortality in swine production^5,6^. This devastation has been observed internationally, as PRRSV-2 gave rise to the “highly pathogenic” (HP) strain that continues to cause mortality in southeast Asia and China^7^. The standard control strategy for PRRSV is to use modified live vaccines that contain an attenuated virus. In the United States there are currently 6 licensed PRRSV-2 vaccine products available^8, 6^. While PRRSV-1 is a potential future concern, there are no currently approved vaccines available, due to the low detection rate and lesser clinical signs presented by this genotype. This study, thus focusses primarily on fitness scoring of PRRSV-2, considering its endemic persistence and high pathogenicity.

PRRSV enters host cells through a multistep process mediated by interactions between viral glycoproteins and cellular receptors on porcine alveolar macrophages (PAMs)^9^. The PRRSV envelope has two main glycoprotein complexes: the GP5-M heterodimer, which is mainly responsible for binding to the host cell receptor, and the GP2-GP3-GP4 trimer, which stabilizes the viral particle and increases its ability to infect cells^10^. The immune response against PRRSV involves B-cell production of antibodies targeting GP5, but viral evasion mechanisms limit neutralization efficacy ^11^. Mutations at key sites in GP5 protein obscure neutralizing epitopes, alters local charge distribution that allows to repel and sterically block antibodies and thereby escape/ become evolutionarily fitter. To this end, we have developed a structurally-guided escape-variant predictor CTRL-V and demonstrated its efficacy in predicting mutations observed in SARS-CoV-2 variants of concern starting with the parental sequence for which the receptor binding domain was co-crystallized with the human ACE2 protein and also commercial Ly-CoV1404 antibodies^12^.

Given the immunodominance of the GP5 glycoprotein in eliciting neutralizing antibody responses, we systematically evaluate the impact of all possible single-point mutations on its antibody escape potential. Anticipating escape tendencies using ML based computational tools is viewed as a promising approach to identify and predict pathogenic variants. EVEscape ^13^ is one such model used for viruses such as SARS-CoV-2, that captures evolutionary landscape for fitness and combines with structural and biophysical viability for a mutant to escape antibody binding by training a variational autoencoder on deep multiple sequence alignments. The integration of protein language models into genomic surveillance pipelines has proven useful for real-time monitoring and characterization of emerging variants^14^. This work builds several folds on EVEscape scoring capabilities to predict critical future variants by replacing the amino acid identity based one hot encoding with state-of-the-art ESM-2^15^ residue embeddings and providing a structural basis of interpreting these variants through receptor interactions. The residue embeddings carry pre-trained evolutionary patterns from millions of protein sequences and even capture spatial relationships between amino acid pairs. This allows us to accurately assess mutational effects to pinpoint functionally important loci for vaccine targets without the need for explicit sequence alignments and experimentally resolved structures.

This computational approach complements experimental efforts that typically assay only a limited subset of PRRSV GP5 variants against available monoclonal antibodies. **Figure 1** shows the porcine host receptor-antibody competitive binding at the *ORF5* receptor binding domain (RBD) of PRRSV transmembrane protein.

**Figure 1:**
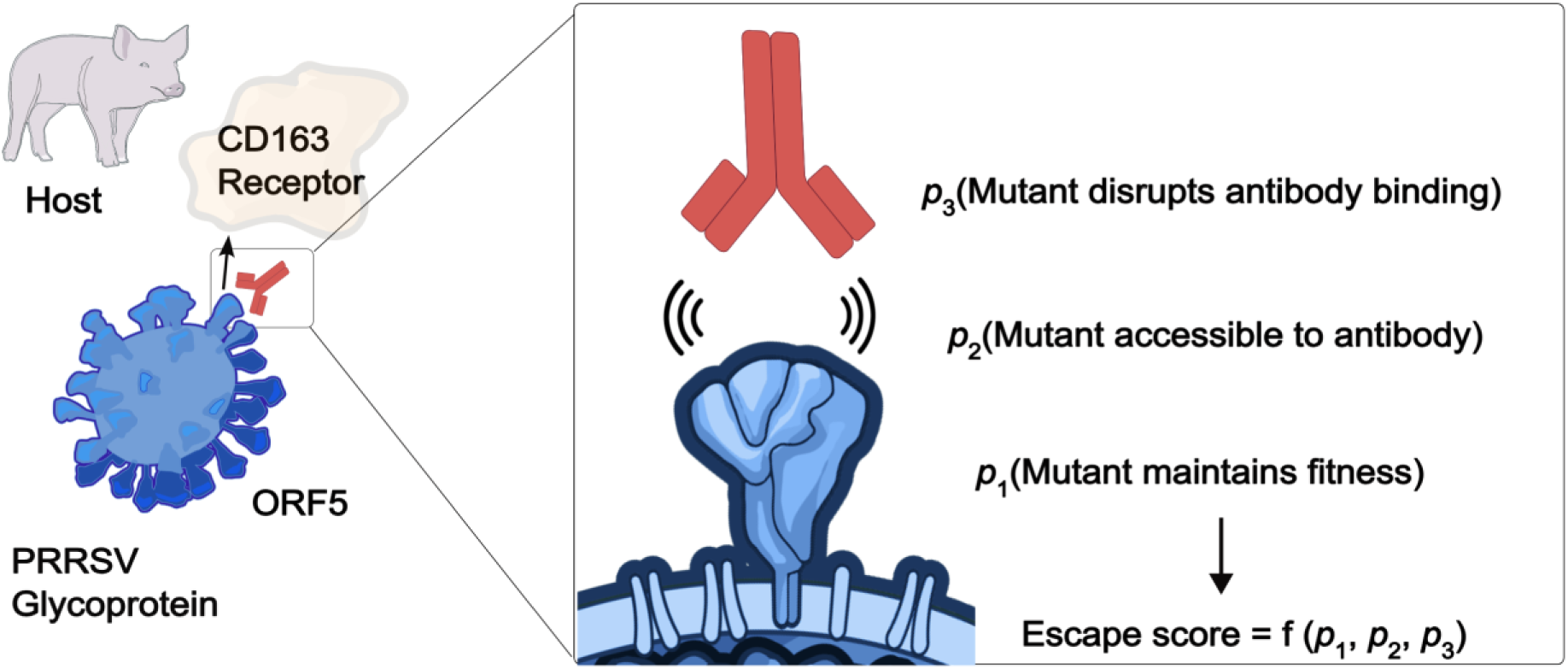
Antibody escape scoring for a single point PRRSV GP5 mutant with EscaPRRS. The genetic construct of GP5 is indicated as ORF5. Illustrations from NIAID NIH BioArt Source^16,17^.

Sequence distributions under natural selection have been previously modelled ^18^ to obey a Boltzmann distribution profile, such that log likelihood of a sequence in its evolutionary landscape is proportional to its fitness, serving as an unbiased zero-shot proxy for mutational tolerance. To calculate the fitness score, the EVE ^19^ model creates one-hot encodings for all residues in an input sequence corresponding to a protein family, which are replaced by ESM-2 embeddings in EscaPRRS. These refined encodings are passed through a Bayesian Variational Autoencoder (VAE). The probabilistic framework revises prior assumptions on the evolutionary latent variables based on the sequence level variability learned by the VAE. To ensure accurate recovery from the latent space, the encoder objective is set to minimize the evidence lower bound (ELBO) with Kullback-Liebler (KL) divergence loss as penalty for deviations from the prior distribution.

The use of 1D convolution in EscaPRRS output captures local dependencies across the sequence positions, preserving the biological realism of the reconstructed sequences, making EscaPRRS more regularized than EVE and generalizable to handle unseen sequences with similar structural motifs. This allows a significant leap over sparse one-hot encodings in EVE, where inputs are positionally independent and outputs are mapped directly to sequence logits. Since ESM-2 embedding representations are sensitive to single point mutations, it supports downstream tasks such as computing evolutionary indices via ELBO estimation and identifying functionally important residues for protein-protein interactions at virus-host interface, while preserving autologous input sequences in the latent space representation.

For an evolutionary fit PRRSV GP5 variant, antibody escape is conferred through structural and physicochemical interactions at the receptor binding domain that allow the viral protein to escape antibody binding while binding to the host receptor. To effectively account for these effects, the surface accessibility conditional to a fit mutant and chemical dissimilarity conditional to effective surface accessibility are incorporated to the escape score.

The accessibility score is taken as the magnitude of the weighted contact number (WCN) for each residue measured as the sum inverse squared distances between all residue pairs in a PDB structure. This way the closer residues to a given residue contribute more to the WCN. For each amino acid residue, the center of mass of the side chain is calculated and all distances are measured between the center of mass coordinates. Fewer neighboring residues empirically indicate that a residue is likely to have more convex surface exposed and accessible to antibody binding. This packing-density dependent surface accessibility/ avidity metric makes it useful to pinpoint obscure epitope regions.

The dissimilarity score, similar to the native EVEscape paradigm is a combined charge-hydrophobicity metric that captures variations in chemical affinity when a residue on a given wild-type structure is substituted with a different one, based on sum of differences in Eisenberg Weiss and charge metrics.

As shown in **Figure 2**, the overall escape ^20^ score is calculated as a product of the fitness, accessibility and dissimilarity scores with suitable logistic temperature scaling factors (*T*_*f*_, *T*_*a*_ and *T*_*d*_), referred to as escape factors as shown in Equation 1.

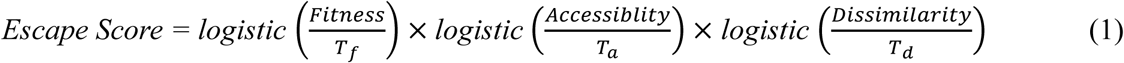

**Figure 2:**
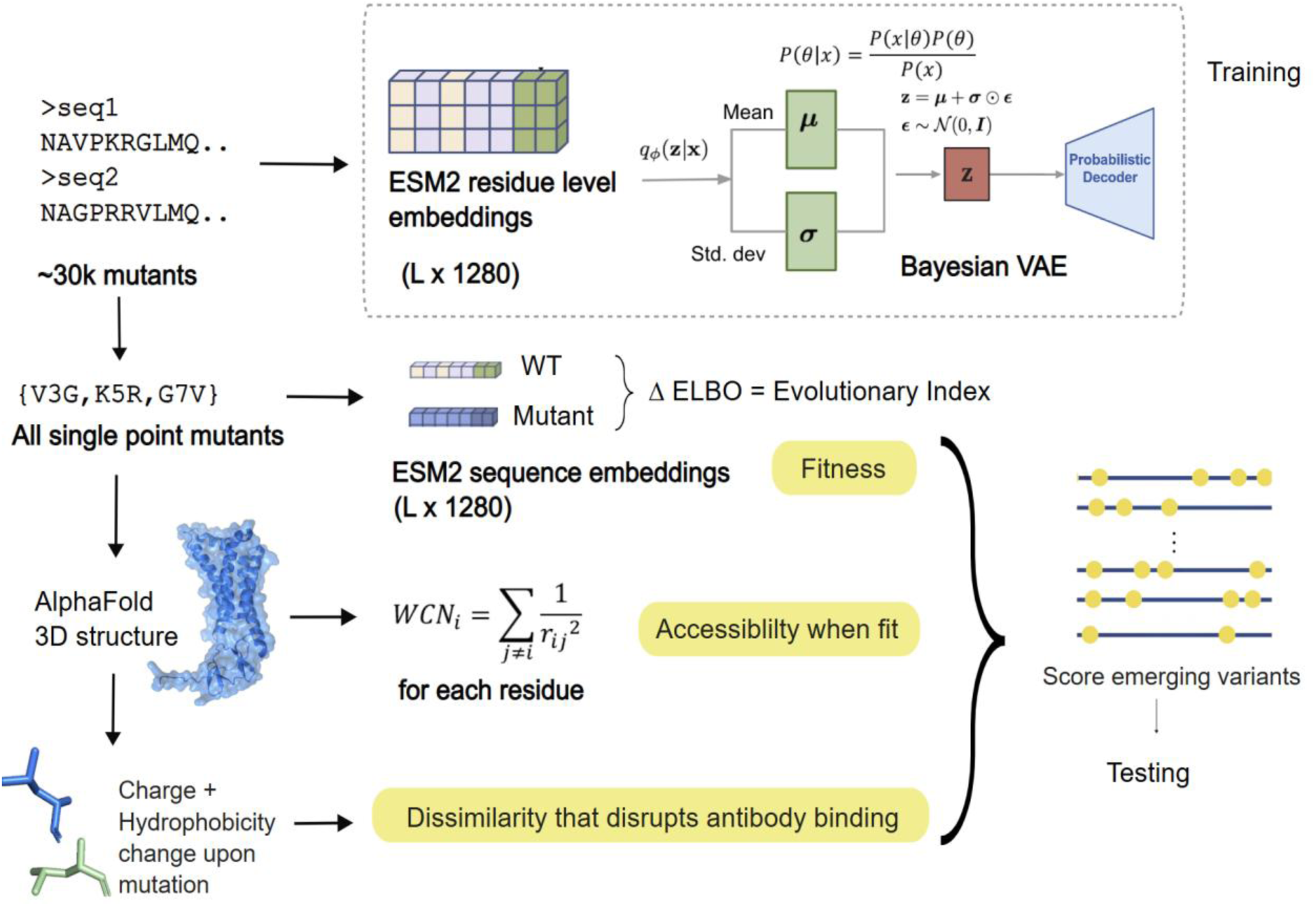
EscaPRRS propensity scoring of emerging sequences incorporating fitness, surface accessibility and residue dissimilarity. ESM-2 embeddings from PRRSV GP5 sequences are trained on a Bayesian Autoencoder that captures variational inference from sequence distributions as in EVE ^19^, to estimate evolutionary fitness of all single point mutants from the wild type. These fitness calculations are enriched with surface avidity and dissimilarity metrics to predict relative escape propensities. The single point mutation effects are aggregated to score emerging variants and sequence constructs for their vulnerability.

In this study, we demonstrate the biological relevance of EscaPRRS antibody escape scores for PRRSV GP5 mutants in capturing seasonal patterns, and providing insights to competitive relative binding affinities at interfacial hotspots for different porcine receptor proteins.

## RESULTS

### Fitness scoring with EscaPRRS

A major component of an escape prediction framework is to demarcate relative fitness of circulating variants as soon as they emerge, to aid parallel efforts on escape-proof vaccine design, as demonstrated in our CTRL-V ^12^ effort. EscaPRRS effectively identified an interpretable fitness parameter and allows two component Gaussian Mixture Model (GMM) based classification used in EVE^19^ model (**Figure 3**, **Supplementary Figure S1**) into benign and pathogenic classes. This structural-basis check helps identify positions where a single point mutation likely compromises protein stability or function over its evolutionary counterparts.

**Figure 3:**
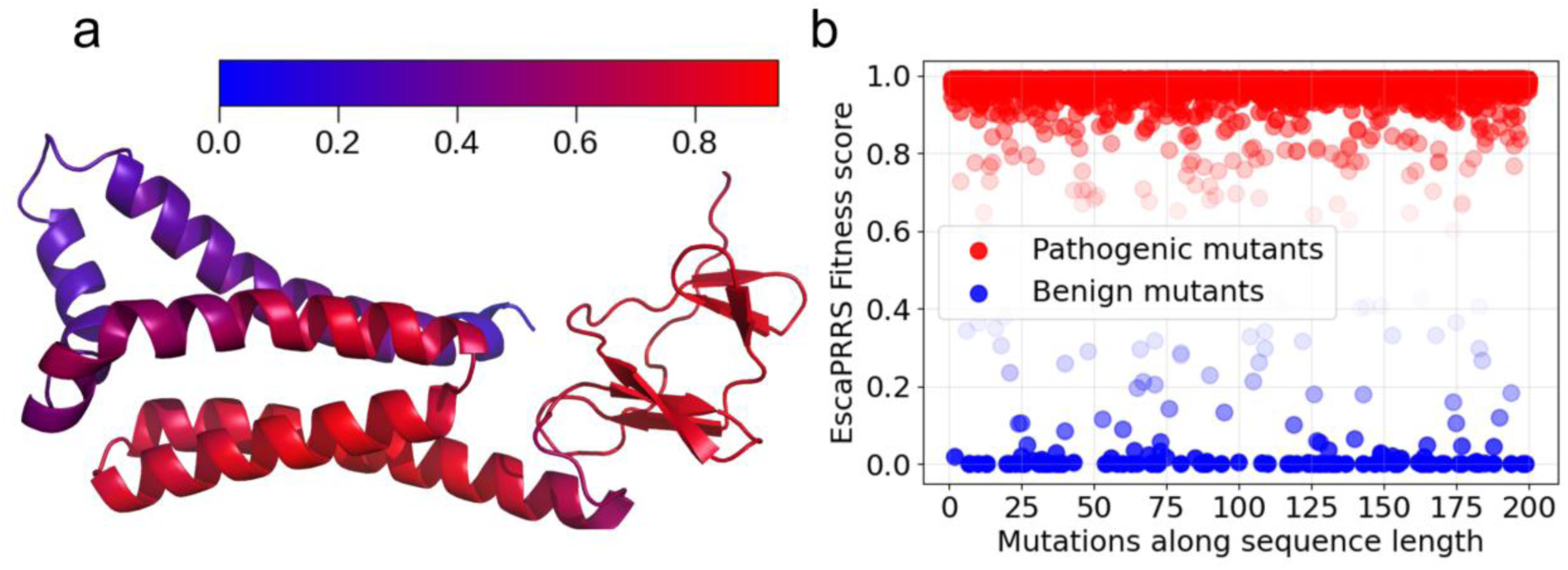
**(a)** Weighted contact number (surface avidity metric) annotated on AlphaFold3 predicted PRRSV *ORF5* GP5 glycoprotein wild-type structure **(b)** Gaussian Mixture Model based classification of the single point mutations in training dataset into benign and pathogenic.

### Ablation analysis

While immune escape mutations come with a fitness cost, the likelihood that a mutation will actually disrupt antibody binding is affected by structural and chemical features at the locus of the mutation. Here, solvent accessibility score captures if a mutation is structurally shielded against exposure to antibody while dissimilarity score measures how much the mutation alters the local charge distribution and chemical properties of the antibody-binding site. These components are transformed into a probability by using a Boltzmann-like partition function (Equation 2), implemented as the SoftMax function in EscaPRRS which helps smoothen the raw embedding data.

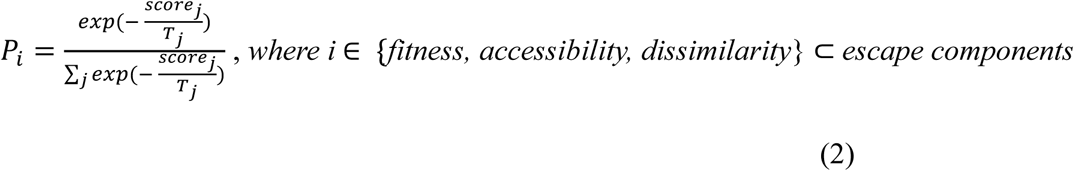

where *T_i_* denotes the temperature (scaling) parameter associated with each escape component *i*, (fitness, accessibility, and dissimilarity) which modulates the sensitivity of the calculated probability to the corresponding *score_i_* that represents the raw score calculated for that component. The denominator ensures the probabilities are normalized to sum to unity.

Each score component is thus scaled by tunable escape factors (*T*_*j*_) to balance the contribution of each component on the final escape propensity, while controlling the randomness of predictions made by the probabilistic model. While low temperature escape factors sharpen the distribution and increase confidence of a prediction, higher escape factor values increase diversity in the output scores. This helps adjust the inherent trade-off between evolutionary plausibility and immune evasion, which is uniquely distributed for different viral proteins based on their structural rigidity and evolutionary plasticity. Noor Youssef *et al.* ^20^ demonstrated hyperparameter searches to optimize escape factors by varying the temperature across a range of values and reported the resulting AUROC values for different viral protein families including flu, HIV, and SARS-Cov-2. Higher AUROC values corresponded to the efficacy of the escape factor setting in the model to predict true escape mutations while avoiding false positives.

Our ablation analysis in **Figure 4**, investigated the determinants of immune escape based on initial escape scores. This analysis revealed that avidity, quantified as structural accessibility reflecting solvent exposure of residues, is the most critical factor in our model. Adjusting the temperature scaling for accessibility resulted in a 38.71% performance decrease, underscoring its strong sensitivity. In contrast, the fitness feature (mutational tolerance) showed a 30.75% increased performance, and dissimilarity (biochemical divergence from wild type) had the smallest effect with 13.66% improvement. Furthermore, the R² score decreased most significantly to 0.1717 upon removal of the accessibility feature, confirming its essential role in PRRSV GP5 mutational escape prediction. The minimal impact observed when removing fitness (R² = 0.9006) and dissimilarity (R² = 0.9042) suggests that structural features predominantly govern escape tendencies, indicating that immune escape mutations are primarily located at structurally accessible sites, likely antibody epitopes, rather than being driven by selection for enhanced receptor binding or altered host mimicry. This is analogous to hypermutability of CDR3 surface exposed loops of antibodies which are constantly under selection pressure to bind to a rapidly evolving antigen ^21,22^.

**Figure 4:**
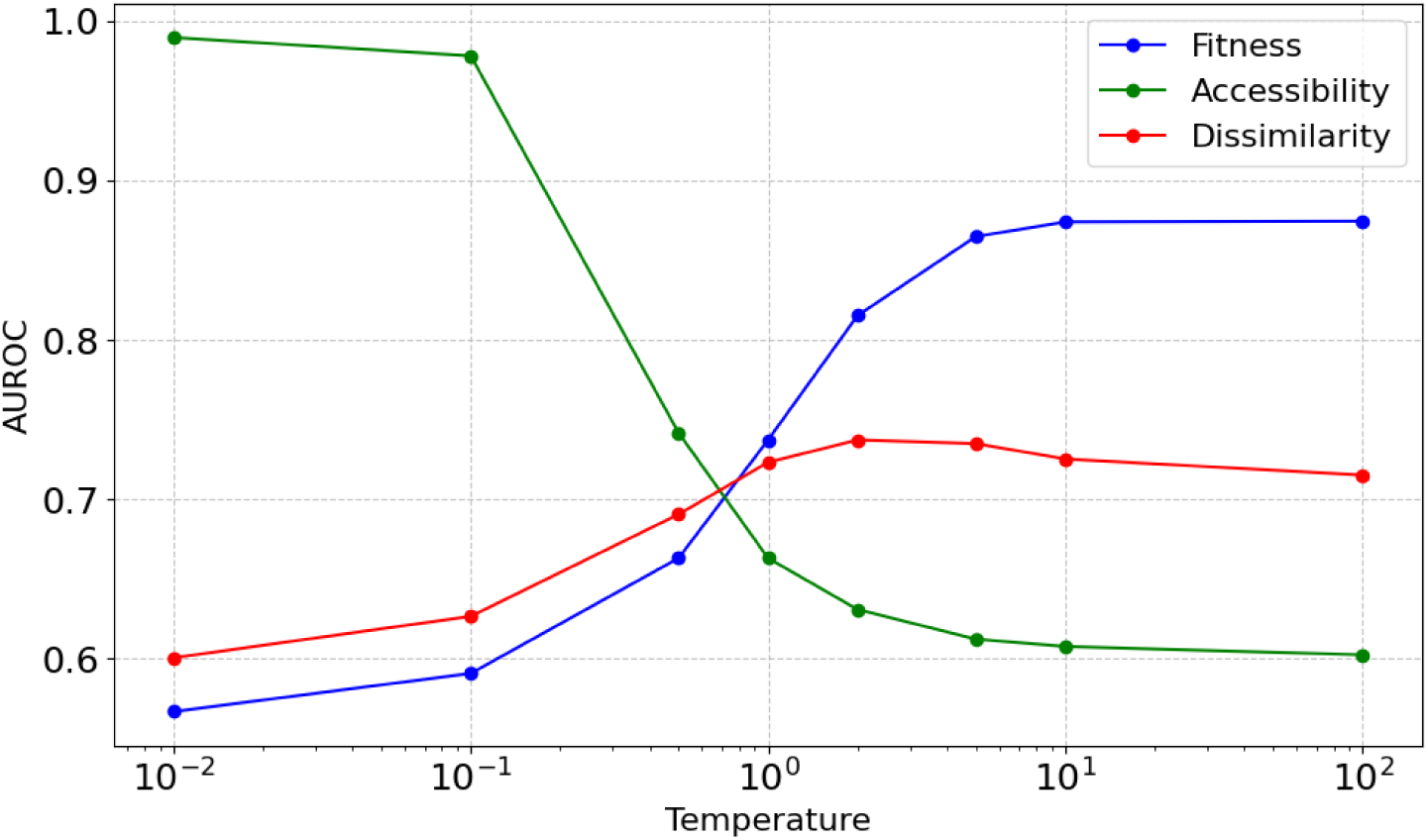
Ablation study showing the impact of removing individual features on the predictive performance of the EscaPRRS based escape scoring model on PRRSV at different temperature values.

On observation of the performance curves for each feature, we see fitness plateaus beyond a scaling of 10, while accessibility feature drops sharply after 0.5 and dissimilarity peaks around 2 to 5. This indicated increasing scaling beyond these regions does not yield meaningful performance gains and could lead to numerical instability. Since the default escape factors (in EVEscape) were initially set to 1,1, and 2, we made a targeted adjustment on escape factors to 5, 0.5 and 5 for fitness, accessibility and dissimilarity respectively to suppress the contribution of surface accessibility on the escape scores for a balanced scoring of mutational escape in EscaPRRS.

### Antibody escape analysis and structural fidelity of EscaPRRS

The EscaPRRS scores calculated with the new escape factors from ablation tests were used to demonstrate the utility of our model in forecasting escape threats from the emerging sequences. **Figure 5** indicates the residues whose mutations consistently relate to higher escape tendencies are conserved at specific positions in the glycoprotein sequence. Analysis of the top 10% EscaPRRS scores among all single-point mutants revealed that the corresponding positions are structurally clustered, indicating spatial proximity within the protein structure.

**Figure 5:**
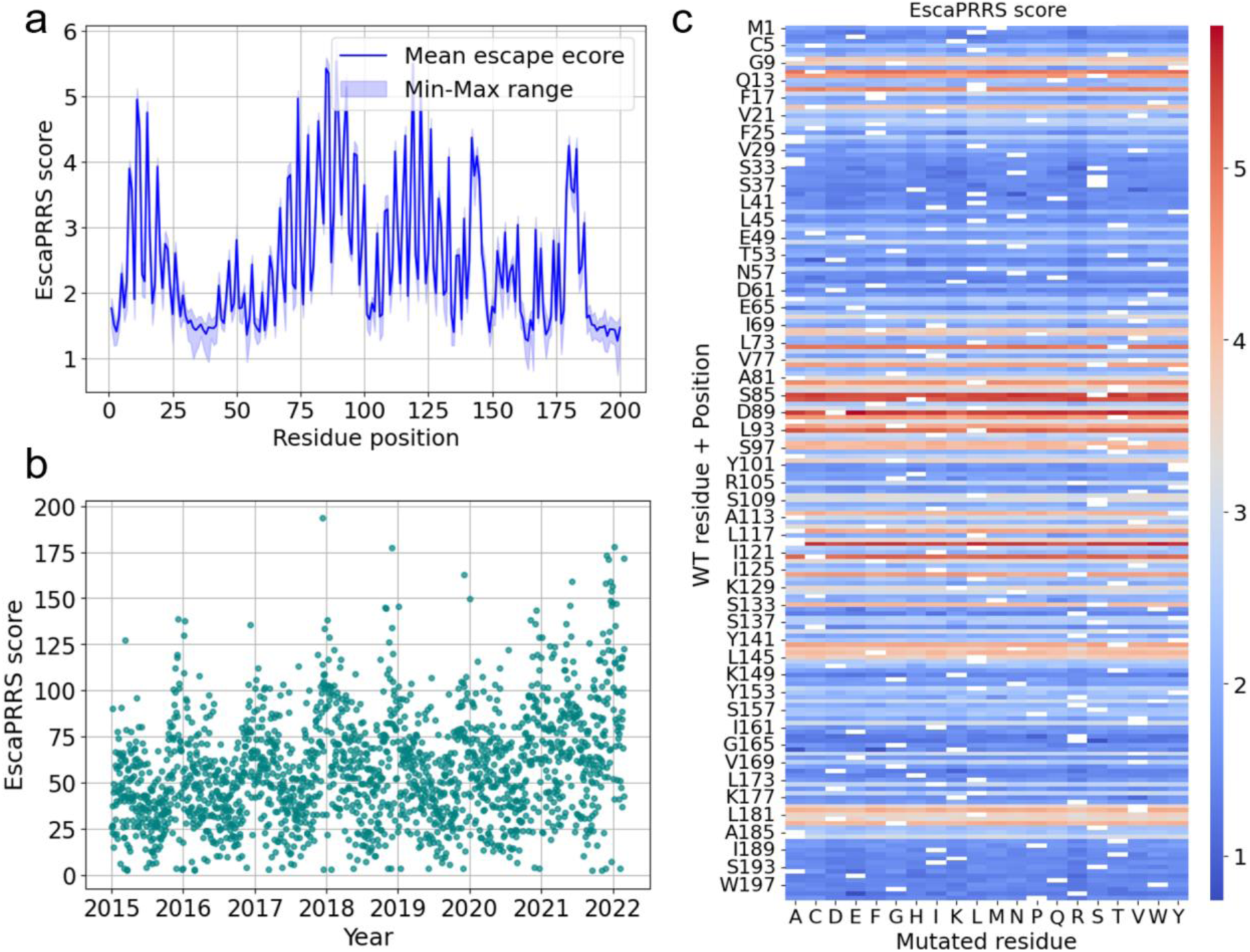
**(a)** Line plot of mean and variable EscaPRRS scores along the sequence length of the mutants **(b)** EscaPRRS scores calculated for 32,536 sequences obtained from Jan 2015 to Feb 2022 (Sequences in the training data).**(c)** Heatmap illustrating the EscaPRRS scores of all possible single mutations to the wild type PRRSV *ORF5* GP5 strain (A detailed heatmap is provided in Supplementary Figure S2).

### Time series forecasting of escape propensities of future variants

To detect general trends in escape tendencies, the EscaPRRS scores (calculated as sigmoid sum of all single point mutation escape scores) for all sequences observed in a given day aggregated to enable time series forecasting.

We use the Prophet model ^23^, developed by Meta, to capture seasonal effects using a flexible Fourier series approach to allow piecewise logistic growth curves, enabling detection and adaptation to sudden changes in regular trends caused by new variants or interventions. Previous studies have shown Prophet’s performance in forecasting outbreaks of infectious diseases and capturing fluctuating incidence patterns to provide early warning signals for rises in pathogenicity ^24–26^.

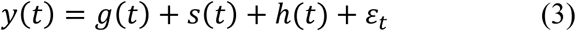

Prophet’s additive model structure as in Equation 3, with trend function g(t), Fourier series seasonal component s(t), holiday effects h(t) and error term ɛ_*t*_ effectively identifies points where trends shift, handling irregularities in sequence data.

With highly diverse lineages observed in emerging PRRSV strains, diagnostic tests on *ORF5* sequences are commonly used to monitor sub-lineage classes and variants in swine populations to track epidemiology. We infer the lineage-wide and variant classification of sequences in our database on PRRSView ^27,28^, which also provides RFLP values and genetic lineage tree for every sequence. This is augmented by occurrence trend data of each variant (classified into accelerating, stable incidence, decelerating, rare/extinct) sourced from PRRSLoom-variants^29,30^. We observe that variants that are reported to be decelerating or extinct correspond to relatively low escape scores, indicating effective antibody neutralization of weaker variants.

The EscaPRRS scores visualized in **Figure 6** indicates a general periodic pattern of higher escape tendencies during the months pertaining to the winter (Nov-Feb) season, suggesting a possibility of higher spread of the virus with higher sequencing efforts.

**Figure 6:**
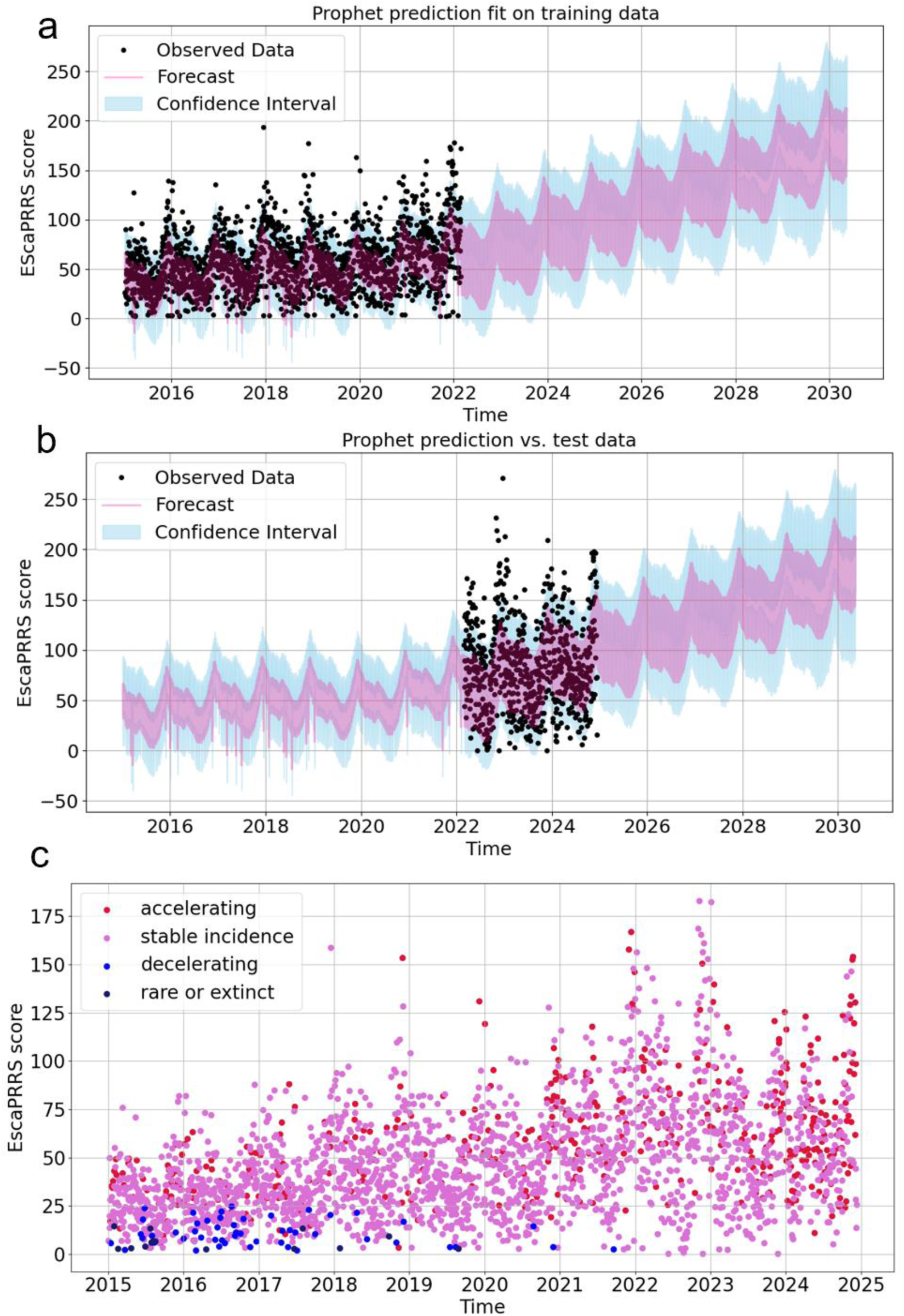
**(a)** Prophet model fit on daily sum EscaPRRS tendencies scored across the training data **(b)** Forecast data on sequences not part of training data. **(c)** Daily sum EscaPRRS scores annotated based on occurrence trends inferred for each variant in 52,622 sequences.

Specific locations where mutations result in increasing (ascending) trend of sum EscaPRRS scores each day and those of descent are mapped onto the GP5 3D structure (**Supplementary Figure S3,S4**). It is observed that the mutations that correspond a descent are localized in a small region with low average escape tendencies, and the descending trend is observed due to the decrease in the number of single point mutations per sequence. We observe an increase in the number of new mutations and missing mutations as we progress through sequences in the descent, while these decrease as we progress through the ascent, which explains the selection pressure towards attaining weaker and stronger variants targeted for antibody escape. The correlation between the number of missing mutations from previous sequences in descent to EscaPRRS scores for current sequences is found to be 0.6069 (Pearson *r*).

To analyze the nature of mutations that are absent in lower EscaPRRS score variants to gauge the significance of structural distribution of mutations on selection pressure towards the weaker variants, we calculate the hydrophobicity change (on the Kyte-Doolittle scale^31^) for each mutation in the list of new and missing mutations corresponding to sequences with an EscaPRRS score of 2.72 to that of 3.13. There is a pronounced increase (9.70) in total hydrophobicity in most missing mutations, while new mutations indicate shift towards lower hydrophobicity (−19.20). The residues at these locations face radially outward **(Supplementary** Figure 5) suggesting weaker mutations being more hydrophobic which could stabilize antibody binding complex at the interfacial hotspots.

To infer time evolution patterns, sequences prior to the training data (recorded before Jan 2015) are recorded and scored for EscaPRRS tendencies (**Supplementary Figure S6**), which reflected trends in sequencing efforts and lineage distributions on the relative vulnerability of the variants towards antibody escape^32^. It is evident that the magnitude of seasonal waves in daily total escape scores along the years followed an overall monotonic increasing trend.

To support our observation of a small fraction of anomalous length sequences in the PRRSV GP5 sequence data, we trained a simple VAE on ESM-2 sequence embeddings to capture indel and insertion patterns. The VAE reconstruction loss is used a proxy to indel probability distribution in an emerging sequence. (**Supplementary Figure S7**).

### Escape tendencies at top receptor binding hotspots

To examine the local effects of single point mutations at interfacial sites of protein-protein interaction between *ORF5* GP5 and the host receptor proteins, we retained the default escape factor weights (1,1 and 2 for fitness, accessibility and dissimilarity) to calculate EscaPRRS scores that highlight the importance of dissimilarity scores given the conditional nature of scoring dissimilarity for a fit mutant with a surface accessible for antibody binding.

Our analysis suggests that GP5 residues that interface with host proteins such as CD163 with higher average EscaPRRS scores across different receptors also tend to exhibit stronger binding affinities, as indicated by HADDOCK3 scores (**Table 1**, **Supplementary Table S1**). This correlation highlights potential antibody target sites. A comparison of binding affinity scores with the maximum site average scores with the interaction residues (**Figure 7**) indicated a Pearson Correlation Coefficient (PCC) of 0.74 with a *p* value of 0.0569. Therefore, this result supports the hypothesis that immune evasion by steric effects, rather than improved receptor usage is the dominant selection pressure driving the evolution and emergence of pathogenic variants.

**Figure 7:**
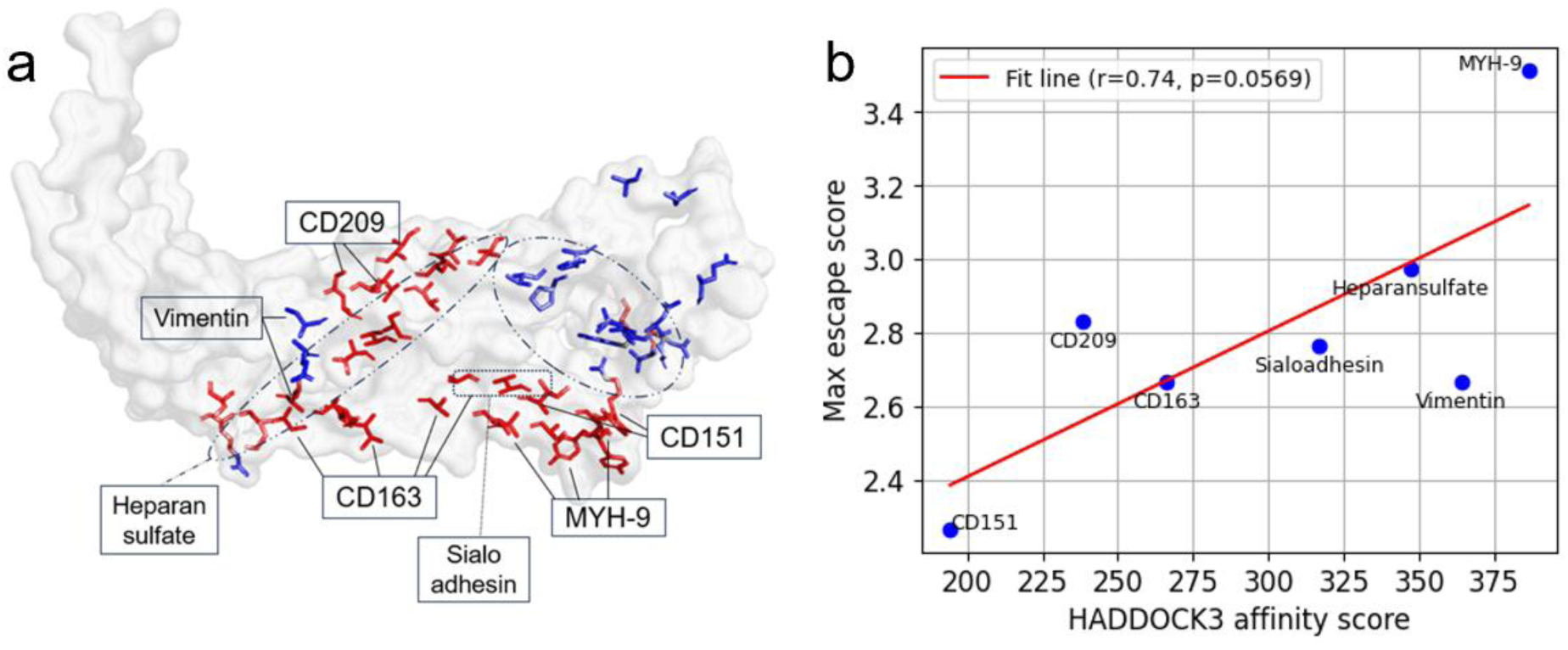
**a)** 3D structure of PRRSV GP5 annotated with residues known to interact to corresponding host receptor proteins. Residue colors indicate EscaPRRS scores where red indicates higher escape propensity and blue corresponds to lower escape propensity. **b)** Scatter plot showing the correlation between HADDOCK3 predicted binding affinity scores and the maximum escape scores among the interfacing GP5 residues with each receptor.

**Table 1:**
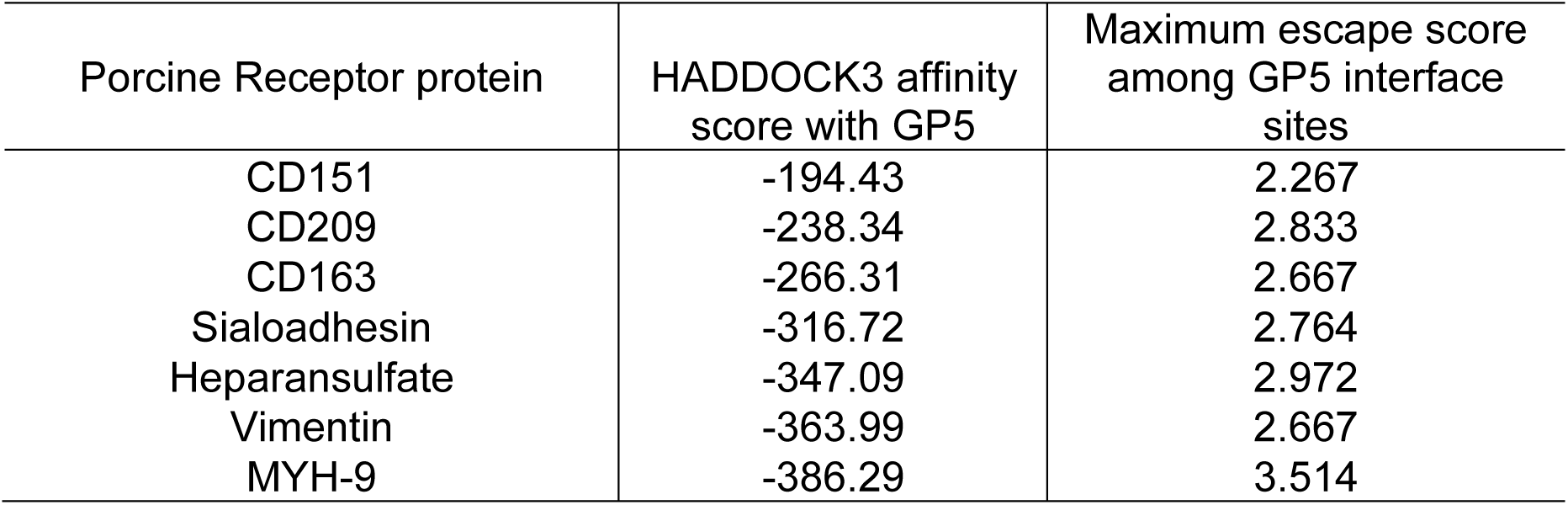
HADDOCK3-predicted affinities of PRRSV *ORF5* glycoprotein GP5 with porcine host proteins CD151, CD209, CD163, Sialoadhesin, Heparansulfate, Vimentin and MYH-9 with the maximum site average EscaPRRS scores among the interfacial residues.

| Porcine Receptor protein | HADDOCK3 affinity score with GP5 | Maximum escape score among GP5 interface sites |
| --- | --- | --- |
| CD151 | -194.43 | 2.267 |
| CD209 | -238.34 | 2.833 |
| CD163 | -266.31 | 2.667 |
| Sialoadhesin | -316.72 | 2.764 |
| Heparansulfate | -347.09 | 2.972 |
| Vimentin | -363.99 | 2.667 |
| MYH-9 | -386.29 | 3.514 |

When a mutant viral protein is deemed evolutionary fit relative to the wild type, it is guided by evolutionary pressure towards improved binding with the host receptor protein, indicated by a higher EscaPRRS fitness score. Given a fit mutant with adequate surface exposed to antibody binding, the antibody-pathogen interaction may still be unstable due to chemical dissimilarity effects at the residue level, causing the viral protein to escape, resulting in higher EscaPRRS scores. Mutants have altered shape complementarity and electrostatic affinity and the receptor binding domain (RBD), leading to the fitness-binding trade-off. An optimal binding affinity that improves viral entry and transmission within the host with lower escape tendencies will allow effective exploration of epitopes as drug targets. ^33^

Our previous work on PRRSV epitopes atlas provides a potential roadmap to expand the scope of EscaPRRS to capture PRRSV variants that infect CD163 deletion variants in pigs^34^. Seq2Bind ^35^, despite being only trained on sequence-pairs and experimental binding affinities, is incredibly structurally aware and shows state-of-the-art performance in recovery of key residues that facilitate binding from each protein, surpassing HADDOCK/ DiffDock and other blind docking tools.

Seq2Bind, at its updated version, is trained on SKEMPI v2 database where pooled features of ESM3^36^ embeddings of two protein sequences are effectively compared to predict interfacial hotspot residues involved in binding and binding affinities. (**Supplementary Table S2, S3**)

To identify representative sequences from the training dataset, we performed t-SNE clustering (**Figure 8**) on ESM-2 embeddings and selected a subset of sequences from distinct clusters, ensuring to cover sequences across all ranges of EscaPRRS score estimates. These sequences were subsequently evaluated for their binding affinity to known porcine proteins using Seq2Bind’s ESM3 model. A moderate correlation was observed between the predicted binding affinities and the EscaPRRS sigmoid sum escape scores for these sequences, with an average Pearson correlation coefficient of r = 0.426, indicating a plausible relationship between antibody escape potential and host receptor binding propensity. Variants with a higher escape propensity with a higher receptor binding affinity is expected to overpower therapeutic interventions that aim to prevent entry to host infection pathways.

**Figure 8:**
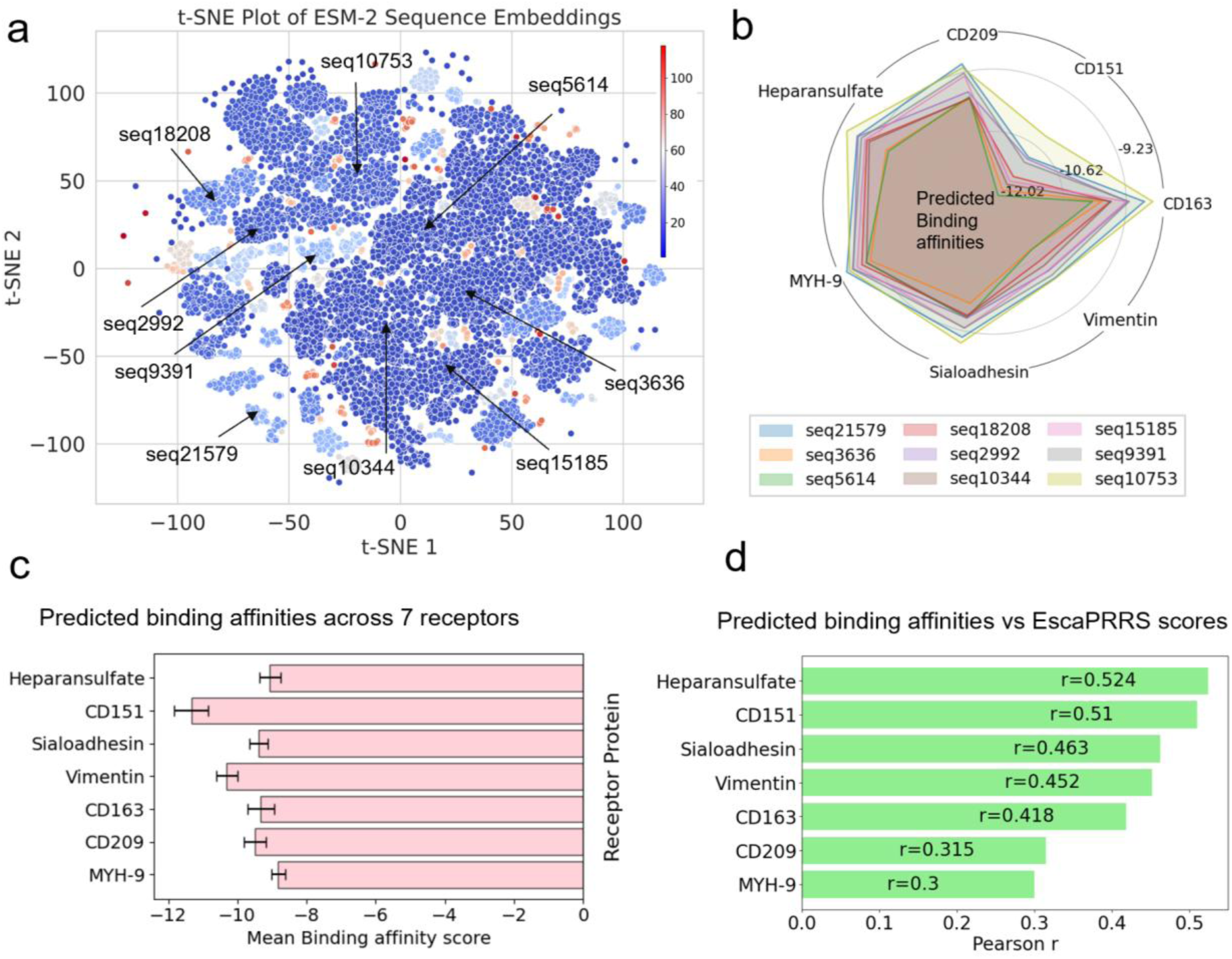
**a)** t-SNE visualization of ESM-2 embeddings derived from 33,717 protein sequences in the training dataset color coded based on DBSCAN cluster numbers. **b)** Radar plot of predicted binding affinities using ESM3 model of Seq2Bind for binding of 7 porcine receptor proteins with GP5 variant sequences chosen across different clusters. **c)** Seq2Bind predicted binding affinity across 7 porcine receptors. Error bars indicate binding score variability across 9 sequences selected across clusters. **d)** Pearson correlation coefficients observed between Seq2Bind predicted binding affinity scores and EscaPRRS scores for 9 sequences across 7 receptor proteins.

However, our Seq2Bind alanine scanning results (**Supplementary Figure S8**) to pinpoint interfacial hotspots for CD163 receptor binding indicate higher binding affinity residues lie in regions with lower site-averaged escape propensities. This supports the fact that mutations that stabilize binding to form spike protein-receptor protein complex may not necessarily ensure that the same mutational changes in the charge distribution at the receptor binding domain (RBD) of PRRSV would confer antibody escape.

Overall, EscaPRRS allows surveillance of PRRSV variants for antibody escape without directly accounting for antibody or receptor information, thus enabling a qualitative check on emerging strains to support escape-aware exploration of vaccine targets. Replacing one hot encodings with protein language model embeddings, which capture a richer evolutionary context, helps address a major limitation in EVEscape. EscaPRRS reduces the reliance on deep multiple sequence alignments for extracting evolutionary patterns and enhances the ability to score the fitness of single point mutations. This also provides a valuable opportunity to assess and predict the escape vulnerability of designed constructs that are antigenically similar to emerging sequences, without heavy dependence on the three-dimensional structures of antigen-antibody complexes. As a result, it becomes possible to identify variants of concern in advance. Aggregating the effects of multiple single point mutations allows exploration of the selection pressures that drive mutants toward favorable binding configurations that facilitate antibody escape, which are inferred from our sequence-based binding prediction tool, Seq2Bind.

Our model can be conveniently modified to handle ESM3 and other protein language model embeddings of any new sequence to score fitness and escape propensities of any viral pathogen. Incorporating dynamic structural features to reflect the complex binding and mechanisms involved in antibody escape presents a valuable direction for future research.

## METHODS

### Modified variational autoencoder (VAE) architecture for PRSSV

We implemented a modified VAE architecture tailored for 200 residue PRRSV GP5 protein sequence modeling using ESM-2 embeddings. The encoder is a multi-layer perceptron (MLP) that flattens the input residue embeddings of shape (200, 1280) and passes them through a series of fully connected layers with ReLU activations and dropout (α = 0.1). The encoder outputs the mean and log-variance vectors for a 50-dimensional latent space.

The decoder is a Bayesian MLP that reconstructs the input from the latent representation. It includes parallel mean and log-variance pathways for each hidden layer, with ReLU activations and dropout. The decoder supports 1D convolution over the output (enabled in our configuration with a depth of 40) and includes a novel temperature scaling mechanism that adjusts the output distribution’s sharpness. Sparsity-inducing mechanisms were disabled in this configuration to prioritize fidelity as ESM embeddings already perform implicit feature selection during training.

This EscaPRRS model is trained using a reconstruction loss (MSE) and a KL divergence term scaled by a configurable factor. This configurable factor ensures balance between reconstruction fidelity and latent space representation.

### Training with ESM embeddings for GP5 sequences of PRRSV

Out of 52,622 experimental PRRSV ORF5 sequences collected between January 2015 and December 2024, 64% were used to train the autoencoder. The remaining sequences from the recent three years were reserved for downstream analysis.

A set of 33,717 nucleotide sequences (timestamped and collated across 7 years) of the PRRSV *ORF5* gene is translated to corresponding amino acid sequences and self-aligned with MMseqs2^37^, with the first sequence being that of the wild type. All sequences are ensured to be of the same lengths and the sequences with anomalous lengths (1181) are removed from the training dataset to avoid introducing hyphenated spaces in the ESM-2 embedding step. The ESM2_t33_650M_UR50D model is used to generate residue-wise (200 × 1280) and average (1 × 1280) embeddings for each sequence.

For each sequence in the input FASTA file, a residue-level embedding is generated using ESM-2. The relative weight of each sequence is computed using a sigmoid-like function as in Equation 4, applied to the cosine similarity between its average embedding and that of the wild-type (reference) sequence.

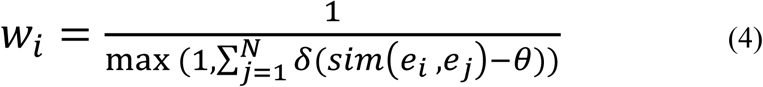

where *e*_*i*_ is the average ESM-2 embedding vector of dimension (1×1280) of a PRRSV GP5 sequence *i*, *sim*(*e*_*i*_, *e*_*j*_) is the cosine similarity of pairwise average embeddings for a similarity threshold *θ*. This allows downweighing sequences with high similarity, promoting uniform distribution and diversity of sequences.

The Bayesian VAE maps the residue-level embeddings onto a 2D latent space to generate mean and variance parameters. The loss values are recorded for 25,000 training steps with a batch size of 256. To compute the evolutionary indices, the VAE-MLP encoder from the original EVE model was modified to accept ESM2 embeddings, which were flattened for latent space representation. These continuous embeddings as regression targets necessitated employing mean square error as the reconstruction loss metric. The evidence lower bound is calculated using Equation 5.

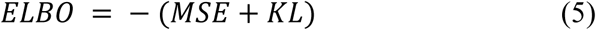

Where MSE is the calculated mean square error for reconstruction of sequences from the latent space representation, and KL is the Kullback-Leibler divergence to penalize sequence probability distribution in the latent space that deviates from the prior distribution in the training data. Summing up these two loss terms allows us to preserve meaningful information during encoding and decoding while encouraging smoothness in the latent space to prevent overfitting of sequence distribution.

A set of all single point mutations in the training data is mapped with the wild type as the reference sequence. For each mutation, the ESM embeddings are generated for the mutant sequence and fed to the trained VAE for ELBO-based evolutionary index calculation, which represents the fitness in the embedding space.

### EscaPRRS scoring

For a set of input sequences, all available single point mutants from the wild type PRRSV GP5 strain are quantified with an evolutionary index, defined as the negative log-likelihood ratio of the mutant sequence relative to the wild type under the trained model. This measures how well a variant aligns with the evolutionary constraints learned from natural sequences. A higher magnitude of an evolutionary index indicates a mutant with high tolerance from pathogenic disruptions. The fitness score is set equal in magnitude to that of the evolutionary index for each mutation.

The structure of wild type PRRSV ORF5 protein is obtained using AlphaFold3 [16] for the translated sequence (GenBank: OR293627.1) from NCBI database. The weighted contact number is calculated for each residue in the wild type structure and is used throughout all sequences towards calculating the overall fitness score for each mutation. The dissimilarity scoring is retained as in the default EVEscape procedure.

For a given single-point mutation, the EscaPRRS score is the product of fitness, solvent accessibility and dissimilarity scores, scaled appropriately with the corresponding escape factors. The escape factors are initially set to 1,1 and 2 for fitness, accessibility and dissimilarity scores respectively based on the default settings in EVEscape model. A subsequent ablation analysis is done to find the optimal scaling factors for the PRRSV GP5 strain. This involved combing through 24 distinct hyperparameter combinations in all.

A list of single point mutations and sites with the top 1% of the predicted EscaPRRS scores are extracted. To score a sequence, the single point mutations in that sequence with respect to the wild type sequences are aggregated to return sum escape score and sigmoid sum escape scores.

### Correlation with Seq2Bind predicted binding affinities

A list of WT sequences for key porcine receptor proteins is sourced from NCBI RefSeq Accession IDs. These include CD163 (XP_020946779.1), Sialoadhesin (XP_020932962.1), CD209 (NP_001123444.1), Heparan Sulfate (XP_020940682.1), Vimentin (XP_005668163.1), CD151 (XP_013845140.1), and MYH-9 (XP_020947197.1). Each of these proteins are parsed with GP5 variants to score the binding affinities, which are compared across models. The predicted affinity scores are evaluated for correlation with EscaPRRS propensity predictions for 9 GP5 variants chosen as representatives from t-SNE clusters of ESM-2 embeddings for sequences in training data.

### Statistical information

Statistical analyses were performed using Microsoft Excel 365’s built-in correlation function and SciPy package in Python. Pairwise relationships between variables were assessed using the Pearson correlation coefficient, which quantifies linear associations between continuous datasets. All tests were two-tailed, evaluating the possibility of both positive and negative correlations. No data transformations were applied prior to analysis. Sample sizes (n) and corresponding correlation values (r) are reported in the figure legends.

## DATA AVAILABILITY

While the training and test sequences used in this work are of proprietary nature, the PRRSView ^27,28^ server shall be referred for a phylogenetic overview of PRRSV-2 *ORF5* GP5 sequences, including BLAST, RFLP scores, lineage and sub-lineage classification tools.

## CODE AVAILABILITY

EscaPRRS is freely accessible at https://agrivax.studio suite of tools and within the StructF tab. Source code to calculate EscaPRRS scores is available at the following Colab link: https://colab.research.google.com/drive/1qs48k65aDkxuxO0rYcCDxtZPHjYlVMrO?usp=sharing. Training and encoder model scripts are available at the following GitHub page: https://github.com/vaishnavey/EscaPRRS

## ACKNOWLEDGEMENTS

This work is partially supported by the Iowa State University Startup Grant (Building a World of Difference Faculty Fellowship), Iowa State University Vice President of Research Seed Grant for PRRSV study, and NSF 22–599, EPSCoR RII Track-1, Award Number DQDBM7FGJPC5 to R.C. V.S.R. acknowledges Dr. Monica H. Lamm, Professor, Department of Chemical and Biological Engineering, Iowa State University for reading and providing feedback on this manuscript.

## AUTHOR CONTRIBUTIONS

R.C. and V.S.R. conceived the idea and did the project design. V.S.R. wrote all the code for the EscaPRRS implementation, performed analyses, and wrote the full first draft of the manuscript. S.D. performed epitope analysis and S.F. worked on understanding and co-opting multiple modules of the EVE model. M.A.Z provided access to the corpus of sequences that are present in PRRSView^27^ through a non-disclosure agreement with R.C. These sequences constitute the training set for EscaPRRS. R.D refined the escape score calculation scripts to set up EscaPRRS tool on AgriVax server. All authors contributed to writing the manuscript.

